# Interpretable gene network inference with nonlinear causality

**DOI:** 10.1101/2025.09.28.678927

**Authors:** Madison S. Krieger, William Gilpin

## Abstract

Interactions among genes orchestrate the growth and survival of all living systems. Such interactions are mediated by vast networks of time-dependent couplings, yet existing strategies for inferring gene networks from omics data do not account for the strong causal constraints acting on nonlinear dynamical systems. As a result, we show here that current gene network inference approaches fail when applied to time series from physical systems. We thus introduce RiCE, a physics-based algorithm for inferring causal interaction graphs using geometric information embedded within time series. We benchmark RiCE against 30 other gene network inference methods, including recent probabilistic machine learning methods, and obtain leading results across 15 datasets spanning diverse experimental modalities and tissue types. RiCE learns physically-interpretable parameters of complex biological systems, requiring low overhead to achieve leading performance. We show diverse applications of RiCE: identifying transition states during the epithelial-mesenchymal transition, classifying cell subtypes during immune cell activation, and determining transcriptional dynamics during endocrinogenesis. Our work demonstrates nonlinearity as a critical inductive bias to accurately infer gene-gene interactions, as well as a key quantitative metric for understanding the dynamic biology of the cell.

---

Understanding causal interactions in omics data is a fundamental challenge in biology, representing a crucial step in explaining how genotype becomes phenotype. Recent developments in high-throughput technologies for quantifying gene expression have led to the creation of many large transcriptomic datasets, making mapping gene-gene causal interaction networks a natural avenue to develop far-reaching computational frameworks. Diverse statistical approaches have been developed to this end, which successfully uncover patterns of co-expression among genes [1]. Yet many interactions among genes are fundamentally nonlinear. For example, a bistable switching mechanism underlies lactose metabolism in *E. coli*, while a pulsed feedback circuit repairs DNA damage [2–5]. Many statistical methods are inappropriate for detecting causality in nonlinear dynamical systems; for instance, nonlinear systems undergo may periods where all variables are strongly correlated and periods where no variables are correlated, making statistical associations a poor tool for causal inference such settings [6]. Ulam’s dictum that “using a term like nonlinear science is like referring to the bulk of zoology as the study of non-elephant animals” is as likely to apply in biological systems as in physics. While some existing approaches address nonlinear interactions indirectly, by leveraging (pseudo)temporal information or inferring unobserved latent variables [1, 7], existing approaches fail to fully leverage the unique structural properties that nonlinear dynamics impose on the dynamical systems underlying gene expression, such as state-dependent stability, bifurcations, and low-dimensional manifolds associated with dynamical coupling. As a result, when leading gene network inference algorithms are applied to time series from physical nonlinear systems, like systems of coupled pendula, they fail to discover physically-meaningful interaction networks (Appendix B).

Here, we introduce **Ri**emannian **C**ausal **E**mbedding (RiCE), a novel physics-based approach to inferring large-scale nonlinear causal relationships from time series. RiCE constructs a model-free local nonlinear approximation of the dynamics of each gene in a time or pseudotime series, and then generalizes classical nonlinear causality measures for dynamical systems to infer directed relationships among genes at scale. We benchmark RiCE against 30 other methods and find that it matches or exceeds state-of-the-art existing network inference benchmarks on real-time expression (DREAM4) and single-cell pseudotime (BEELINE and McCalla), with its advantage growing as the underlying systems become more strongly nonlinear. As a physics-based model, RiCE is highly scalable and produces interpretable internal representations, which we illustrate by identifying cells entering transition states during the epithelial-mesenchymal transition, and by probing transcriptional dynamics during endocrinogenesis. On a large-scale immune cell dataset, RiCE discovers a network that captures known biological markers like highly-connected transcription factors, and subcommunities associated with immune cell subtypes.

## I. APPROACH

We generalize nonlinear causality detection, a class of physics-based methods originally developed to infer interspecies interactions from ecological time series [6]. Under this framework, a one-way causal interaction exists between two genes if information from upstream gene incrementally improves forecasts of the downstream time series.

Our approach is outlined in Figure 1. For each gene’s time series, we use time delay embedding to estimate the local manifold of the dynamical system underlying the gene’s dynamics. For each point on each reconstructed manifold, we estimate the local stability matrix of the dynamics, assigning each timepoint a nonlinearity score corresponding to the local curvature of the dynamical manifold, indicating deviation from linearity. We use this matrix to weight the neighboring timepoints on the manifold, and then train a linear self-forecast model for each gene. We then select a single target gene’s dynamical manifold, and train a ridge regression model to forecast that gene’s dynamics, using all other genes’ forecast models as covariates. The relative regression weight of each upstream gene indicates its causal influence. By training the full model end-to-end, we obtain (1) A *nonlinearity score* derived from the local stability matrix of each gene-timepoint, and (2) an *interaction matrix* representing the relative contribution of each gene to improving each other gene’s forecast. We repeat our procedure for downsampled versions of all time series, and downweight interactions that fail to strengthen with additional timepoints, indicating that the assumption of an underlying dynamical system is weak for that particular pair of variables. We optionally prune indirect interactions, in which two variables have an apparent causal link mediated by an intermediate variable. When applied to deterministic time series from physical dynamical systems, RiCE recovers directed causal relationships, in contrast to standard network inference techniques (Appendix B) We describe our approach mathematically in Appendix C, and we describe its relationship with other empirical causality methods in Appendix D.

**Figure 1.**
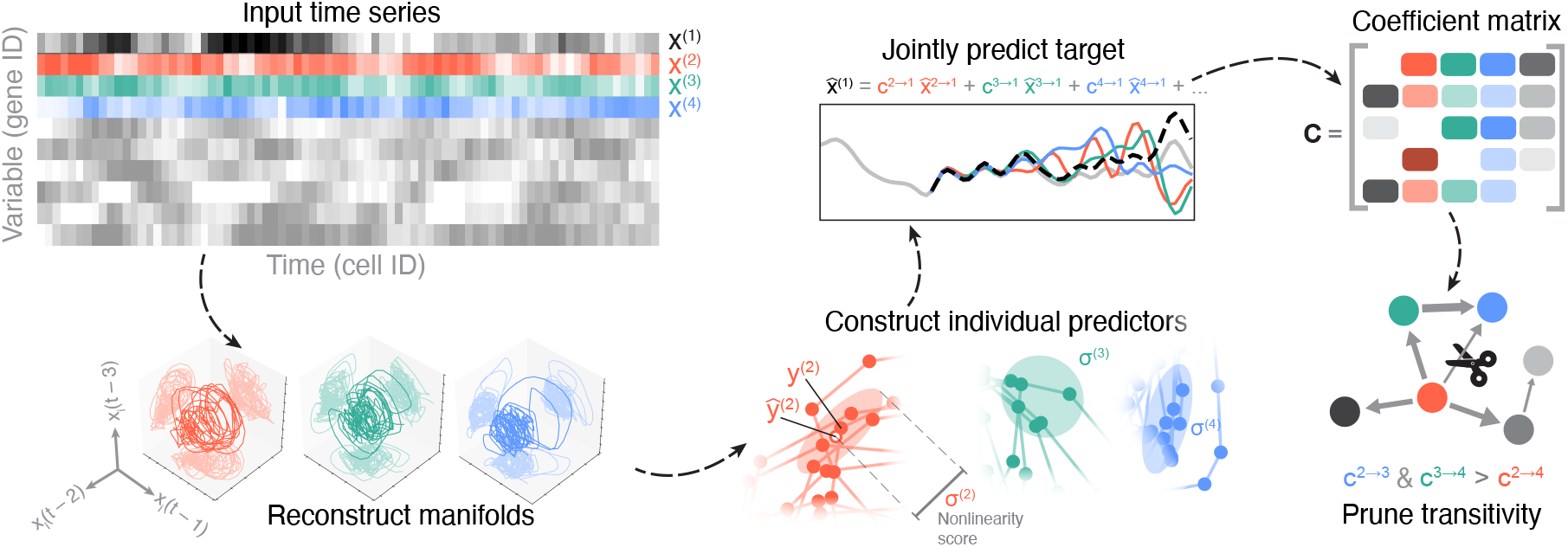
RiCE uses nonlinear causality to infer gene networks. A set of univariate time series are lifted to multivariate dynamical manifolds using time-delay embedding. A locally-weighted forecast model is separately fit to each multivariate time series, by fitting the local stability matrix (Jacobian) of a nonlinear dynamical system. An ensemble of these models is then used to predict each time series, and the relative weights produce entries in the causal connectivity matrix. The resulting network is optionally pruned for indirect interactions.

## II. RESULTS

### A. RiCE discovers gene interaction networks from time series

We evaluate RiCE’s empirical performance relative to other network inference techniques across a range of datasets. We consider 30 reference methods, spanning classical statistical inference techniques to state-of-the-art algorithms specialized for network inference.

#### Benchmark methods

Our benchmark models include: (1) Statistical or information theoretic relationships among genes (MutInfo, PCorr, PIDC, CLR, ARACNE, BayesRidge) and regression models (GLASSO, AdaBoost, GRNBoost2, GENIE3, TIGRESS), which do not directly use temporal information. (2) Temporal models that either directly model an underlying dynamical system (SCODE, dynGENIE3), or use temporal causality measures (SWING-RF). (3) Non-parametric statistical models based on artificial neural networks, which recently have shown leading results for single-cell pseudotime data (RegDiff, DEEPSEM). We further describe all benchmark methods in Appendix G. We also benchmark ablations of RiCE with different components removed. *NoCycles* prunes potential transitive interactions to produce a network with a traditional top-down directed regulatory structure, *NoEnsemble* separately analyzes each pair of genes and thus removes any combined causal driving, *S-map* removes both ensembling and adaptive reweighting of nonlinear neighbors; this ablation’s name is derived from its similarity to a prior method for measuring nonlinearity in time series [8]. *CCM* removes ensembling, adaptive weighting, and replaces all least-squares fits with distance-weighted averages over neighbors, corresponding to an earlier non-linear causality method [6]. To ensure consistency, we re-run and score all benchmark models, using the original authors’ code when available. We do not provide transcription factor labels or other annotations to any methods. To ensure that all benchmark methods function correctly, we test all implementations on a trivial dataset comprising white noise time series with known correlation structure, to verify that their performance scales with interaction strength Appendix G. We do not tune any hyperparameters for RiCE; we use single-shot calculation at fixed embedding dimension (*d*_*E*_ = 3, the minimal embedding required to resolve aperiodic dynamics).

#### Datasets

We first perform all benchmarks on the DREAM4 100-gene challenge. This dataset consists of networks subsampled from the *E. coli* and *S. Cerevisae* gene regulatory networks, with transcriptional dynamics simulated using a stochastic kinetic model. To test to role of nonlinearity, we extend DREAM4 by introducing a new dataset, *Twist*. This corresponds to the original 100-gene DREAM4 networks, as well as 30 new yeast or gene subnetworks randomly-sampled using the same greedy heuristic as the original work [9]. However, we replace the linear kinetic model with a nonlinear differential equation describing coupled mRNA-protein dynamics. We consider two large-scale single-cell pseudotime datasets (BEELINE and McCalla) in Section II B.

#### Metrics

We report accuracy as the Area Under the Precision-Recall Curve (AUPRC), a metric that is robust to class imbalance due to network sparsity. However, we find similar results with two other standard metrics: Area Under the Receiver-Operator Characteristic (AUROC) and Early Precision Ratio (EPR) (Appendix G).

#### Benchmark results

RiCE performs competitively with top methods across all benchmark datasets, with performance increasing with the degree of induced or measured nonlinearity. On DREAM4, the average performance of RiCE matches dynGENIE3, the best-performing alternative method in our experiments; however, we note that the small scale of DREAM4 implies a degree of performance saturation [1]. On *Twist*, a dataset with strong intrinsic nonlinearity, the performance of RiCE strongly exceeds the other methods (Fig. 2A). To better understand this effect, we regenerate the Twist dataset with varying degrees of intrinsic nonlinearity (with DREAM4 representing zero nonlinearity), and we find that only RiCE and its ablations retain performance (Fig. 2C). We thus conclude that nonlinearity produces cryptic interactions, challenging existing network inference approaches that lack explicit inductive biases for nonlinear time series.

**Figure 2.**
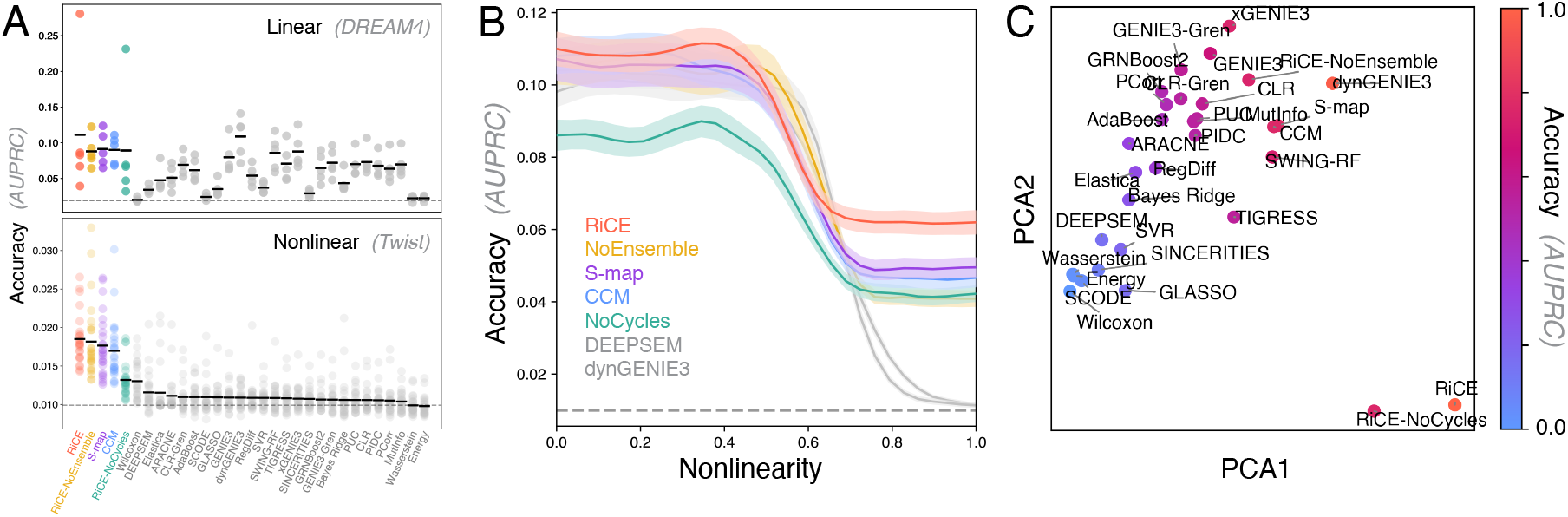
RiCE effectively identifies nonlinear interactions in gene interaction networks. (A). Accuracy of network link identification (AUPRC) across RiCE, ablations of RiCE’s key components, and benchmark approaches. All methods are applied to a standard benchmark dataset of linear stochastic gene regulatory dynamics (top, the DREAM4 100-gene challenge), and a version of this dataset modified to exhibit intrinsic nonlinear interactions (bottom, Twist). The expected AUPRC of a random classifier is indicated by a dashed line on the graphs. (B). Relative performance of RiCE and other dynamics-based inference algorithms, as the degree of nonlinear dynamics in the underlying dataset increases. The best-performing methods on DREAM4 (Linear) are shown for comparison. (C). The prediction accuracy of all benchmark models across all DREAM4-100 datasets, projected onto the first two principal components across all datasets.

To confirm that RiCE functions differently than other methods, we take individual AUPRC scores for each benchmark method across DREAM4-100, and treat these scores as featurizations of each model. In Fig. 2C we project all models onto the first two principal components. We find that RiCE quantitatively differs from other methods: one principal component indicates average performance, but second isolates RiCE from other methods. Statistical and information-theoretic methods, such as GLASSO or Mutual Information, cluster together along this axis, while RiCE and its ablations, along with time-delay methods like SWING-RF or dynGENIE3, cluster separately, implying that this axis distinguishes methods that identify dynamical features.

### B. RiCE infers regulatory networks from single-cell pseudotime

We next evaluate RiCE on two large-scale benchmark single-cell RNAseq datasets, BEELINE and McCalla [7, 10]. BEELINE consists of datasets of human embryonic stem cells (hESC), mouse bone-marrow-derived dendritic cells (mDC), mouse embryonic stem cells (mESC), human mature hepatocytes (hHEP), and three lineages of mouse hematopoietic stem cells (mHSC-E, mHSC-L, mHSC-GM). For this dataset we use as a gold standard a protein interaction network (STRING) curated by the benchmark authors [11]. McCalla consists of two yeast perturbation experiments (yeastA2S, yeastFBS), one human embryonic stem cell differentiation time course (hESC), mouse dendritic cell response (mDC), and a mouse reprogramming dataset (mESC). For McCalla, we use a gold standard dataset curated by the original authors consisting of a union of knockdown and ChiP-seq measured interactions [11]. To test the effect of network size, for both datasets we reconstruct net-works among the 500- and 1000 highest-variance genes, though for BEELINE we force the inclusion of all transcription factors to match prior work [10]. We order the cells in both datasets by pseudotime [12].

We find that RiCE matches or exceeds the performance of other methods on both benchmarks (Fig. S4A), competing with recent artificial neural network approaches that produce leading results on both datasets. This performance is noteworthy, because RiCE is a model-free dynamical systems tool without particular specialization for omics data. As a result, we find that RiCE requires less computational resources relative to its performance than other methods (Fig. S5).

As with any performance benchmark, network inference methods exhibit a “no free lunch” tradeoff between generality and accuracy on a given data class [13]. We therefore interpret RiCE’s performance as evidence that nonlinearity comprises a strong inductive bias for gene interaction networks encountered in many single-cell experiments. To confirm this effect, we quantify the intrinsic nonlinearity of each dataset, based on the average magnitude of the RiCE nonlinearity score across all pseudotimepoints (Fig. S4B). We find that more intrinsically nonlinear datasets produce stronger performance with RiCE, indicating association between a dynamics-level property and the suitability of physics-based causal approaches.

### C. RiCE identifies transition states during the epithelial-to-mesenchymal transition

The nonlinearity score internally produced by RiCE quantifies the time-dependent nonlinearity of the underlying dynamical system. We next consider how the RiCE nonlinearity score reveals characteristic dynamical states across cells. We consider a dataset of squamous cell carcinoma in mouse during the epithelial-to-mesenchymal transition (EMT), with cells ordered by pseudotime. This Smart-Seq2 dataset comprises 295 cells and 1408 genes after accounting for quality and count depth [14].

On the chaotic Lorenz attractor (Fig. 4A, Fig. S2), the dynamics traverse nearly-linear metastable cycles near lobes of the attractor, leading to low nonlinearity score. However, the RiCE nonlinearity score increases in the transition region between lobes, a highly-unstable region responsible for the overall chaotic dynamics of the system [15]. Visualizing the RiCE nonlinearity scores of cells in the EMT dataset reveals a collection of highly-nonlinear cells (Fig. 4A), which occur at intermediate pseudotimes across the transition (Fig. 4B). We observe the same effect when directly comparing nonlinearity versus the expression level of *COL3A1*, a pure epithelial marker[16], underscoring that the RiCE nonlinearity score does not require global pseudotime (Fig. 4C).

**Figure 3.**
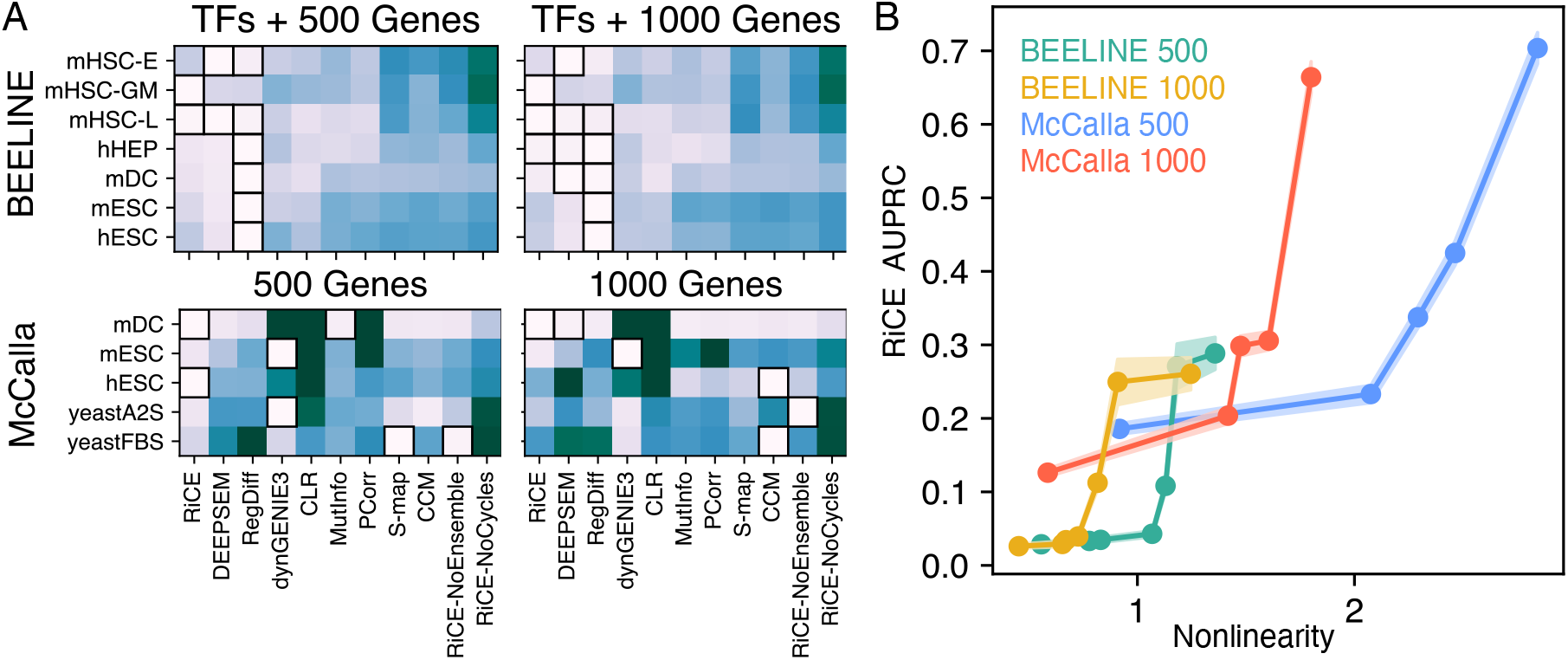
RiCE discovers regulatory interactions from single-cell pseudotime. (A) The accuracy (AUPRC) of RiCE relative to the best-performing benchmark models on the BEELINE and McCalla single-cell gene expression benchmarks. Experimentally-validated gold standard networks (STRING protein interactions for BEELINE, ChIP-seq plus perturbations for McCalla) are used as ground-truth networks. Boxes indicate highest-performing methods for each dataset within one standard error across bootstrapped replicates (resampled cells). (B) Accuracy versus the intrinsic nonlinearity of each dataset, estimated by averaging the RiCE nonlinearity score across all cells. Shaded regions are standard errors across bootstraps.

**Figure 4.**
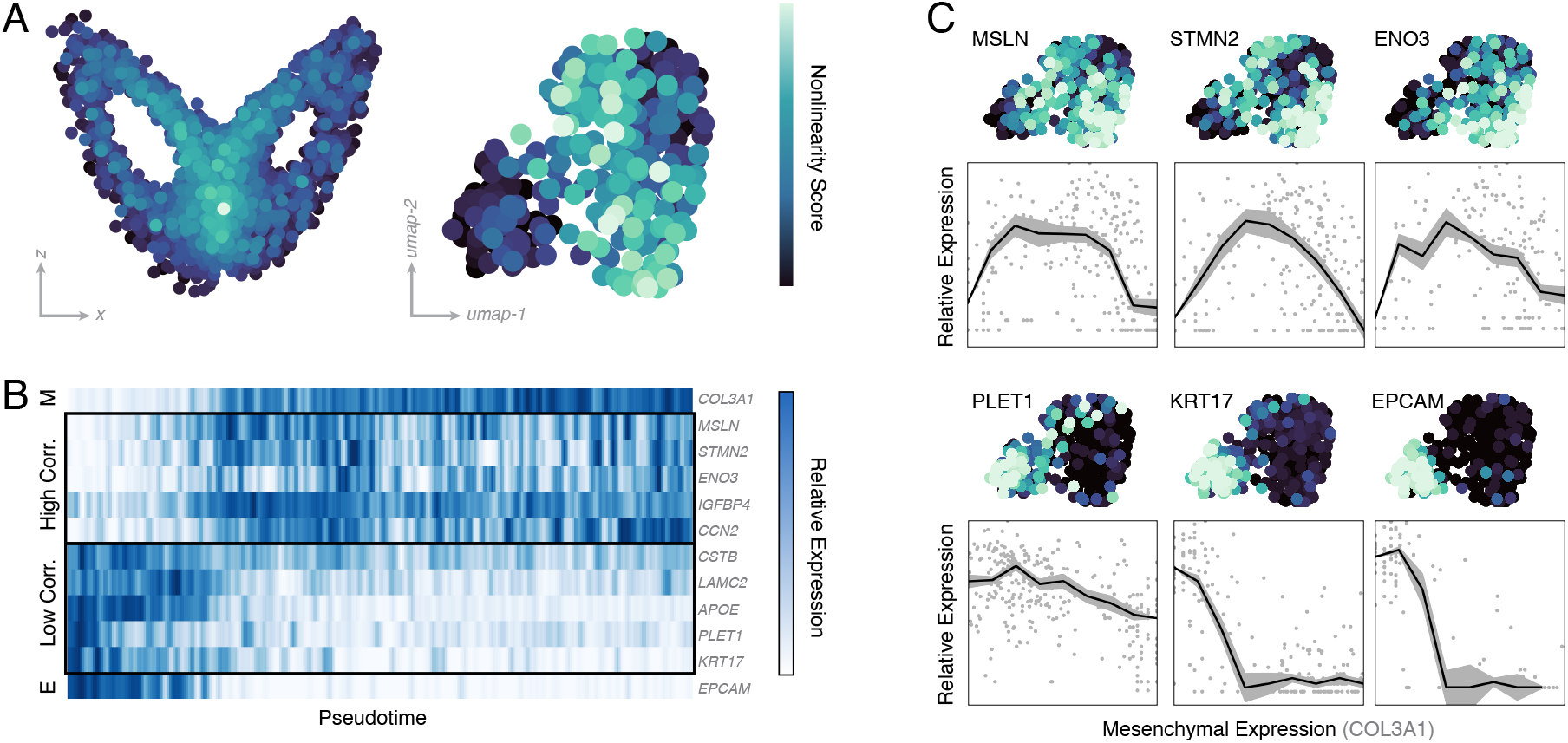
RiCE identifies transition states during the epithelial-mesenchymal transition (EMT). (A). Calculation of the RiCE nonlinearity score on a reference chaotic dynamical system (the Lorenz attractor), and on a dataset of 295 mouse cells undergoing EMT. (B) Expression level versus pseudotime for an epithelial state marker gene (E), the five genes most correlated with the RiCE nonlinearity score, the five genes least correlated, and an epithelial marker gene (M). (C) Expression levels of the most (top) and least (bottom) nonlinear genes, versus the expression level of the pure mesenchymal marker *COL3A1*. Lines are medians of evenly-spaced bins, and ranges are standard errors across 10^3^ bootstrapped replicates.

The proteins associated with the top three genes (Mesothelin, Stathmin-2, Enolase) all have previously been shown to regulate EMT in knockdown experiments [17–19]. In contrast, molecules associated with the lowest-scoring genes (Placenta Expressed Transcript 1, keratin, and EPCAM) exhibit monotonic profiles, and correspond primarily to structural molecules used as markers of either pure epithelial or mesenchymal states [14, 20]. The RiCE nonlinearity score therefore correlates with instability and decision states, and inversely correlates with fate commitment.

### D. RiCE identifies gene subnetworks driving immune cell differentiation

We use RiCE to estimate gene networks from a collection of 16,000 peripheral blood mononuclear cells (PBMC) from a single healthy donor (10x Genomics 3’ v3.1, Chromium X). We subselect CD4+ and CD8+ cells based on an annotated reference, retaining the 3500 highest-variance genes after quality control [22]. We apply RiCE, and find that the resulting network exhibits hierarchical structure, with the ten highest outdegree genes among the 3500 nodes including two canonical transcription factors associated with immune cell differentiation (PRDM2, XBP1), as well as three immune-specific genes (ZAP70, SAT1, COMMD7) [23]. We next perform multilevel community detection on the RiCE causal graph using modularity clustering with varying resolution, resulting in a dendrogram relating communities at varying resolutions (Fig. 5B). We generate community labels by testing for enrichment relative to cell types in the Azimuth PMC reference ontology generated using cell surface markers (CITE-seq) [21]. We consider only enriched types with statistical significance (Benjamini-Hochberg adjusted p-value less than 0.05) [24]. At the coarsest granularity, we find a clear distinction between a gene network associated with B-cells versus T-cells (Fig. 5B). We note that the cell population itself contains only CD8/CD4 T-cells, likely due to overlapping functions and phenotypes between T cells and B cells such as IL-10 secretion by regulatory subsets [25] As we increase the granularity of community detection, we isolate a state enriched for proliferation (SOX and other genes), consistent with intermediates in the immune cell activation hierarchy. We assign each community parents by testing for statistical enrichment of their constituent genes relative to each parent’s gene set. The resulting dendrogram recapitulates the major stages of immune cell activation (Fig. 5B) [25].

**Figure 5.**
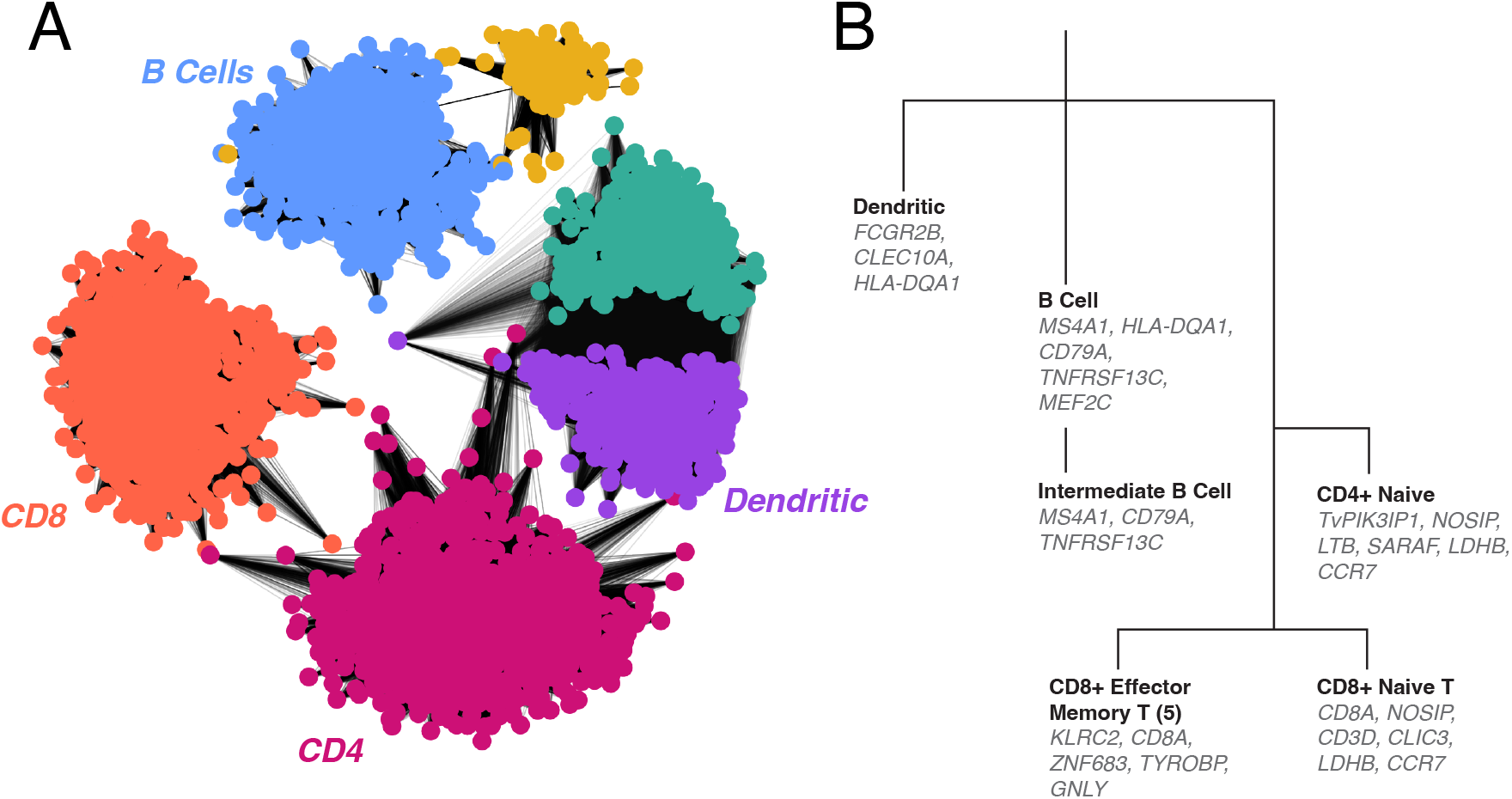
A RiCE-based bioinformatics analysis of immune cells predicts cell-type specific gene networks. (A) The gene-gene causal networks identified by RiCE from peripheral blood mononuclear cells (PBMC) from a single donor. Colors correspond to communities detected via unsupervised modularity clustering on the graph, and labels are generated by testing for enrichment of genes relative to reference PBMC ontology (Azimuth) [21]. The network visualization uses a community-aware force directed layout. (B) A dendrogram generated by varying the resolution of the modularity clustering algorithm, and then assigning finer clusters to parent groups based on significant enrichment. Labels and indicative genes are generated via enrichment relative to the reference ontology.

### E. RiCE identifies transcriptional dynamics

Because RiCE implicitly fits a continuous nonlinear system across timepoints, we next consider whether this flow field agrees with the experimentally-measured transcriptional dynamics. We consider a dataset comprising the 2000 highest-varying genes from an experiment of 3696 cells during pancreatic endocrinogenesis [26]. Prior RNA velocity analysis of this dataset reveals rich transcriptional dynamics, including diverging regions associated with cell differentiation, and vortical regions associated with cycling endocrine progenitors [27].

We find that a cell’s nonlinearity score correlates with both the speed (magnitude) and the divergence of the RNA velocity vector field (Fig. 6A). In dynamical systems, the nonlinearity of a dissipative system dictates its instantaneous phase space contraction rate (the Lyapunov exponents). Across all cells, we find correlation of 0.23 ± 0.02 between the RiCE nonlinearity score and the divergence of the RNA velocity field, consistent with non-linearity occurring in dissipative regions of phase space. We find a correlation of *−*0.26 ± 0.03 between the RiCE nonlinearity score and overall RNA speed, indicating that cells exhibit more linear dynamics when transcriptional dynamics proceed more rapidly. Both results are consistent with the expected behavior of these quantities for linear and nonlinear dynamical systems (Appendix E). Comparing the cell-wise nonlinearity with the latent time inferred from RNA Velocity, we observe that nonlinearity peaks early during the endocrinogenesis cycle, a period of strong differentiation. The five most nonlinear individual genes (SPARC, RPS8, RPS4X, RPS19, RPL32) all significantly overlap with secretory and endothelial programs when compared against the broad Tabula Sapiens reference ontology (Benjamini-Hochberg adjusted p-value *<* 10^−5^) [24, 28]. Conversely, the five least non-linear genes (CPE, PCSK1N, FAM183B, RBP4, SCG) include marker genes for terminal fates, and enrich for terminal pancreatic cell types when compared to tissue-specific gene ontologies (Adjusted p-value *<* 10^−3^). We thus find that more linear dynamics emerge after fate commitment. These results align with the expected behavior of a Waddington landscape model, in which cells exhibit locally-linear dynamics near fixed points associated with terminal fates, and nonlinearity on boundaries between fates [29]. Our findings also align with empirical results showing this effect in endocrinogenesis [30–32].

**Figure 6.**
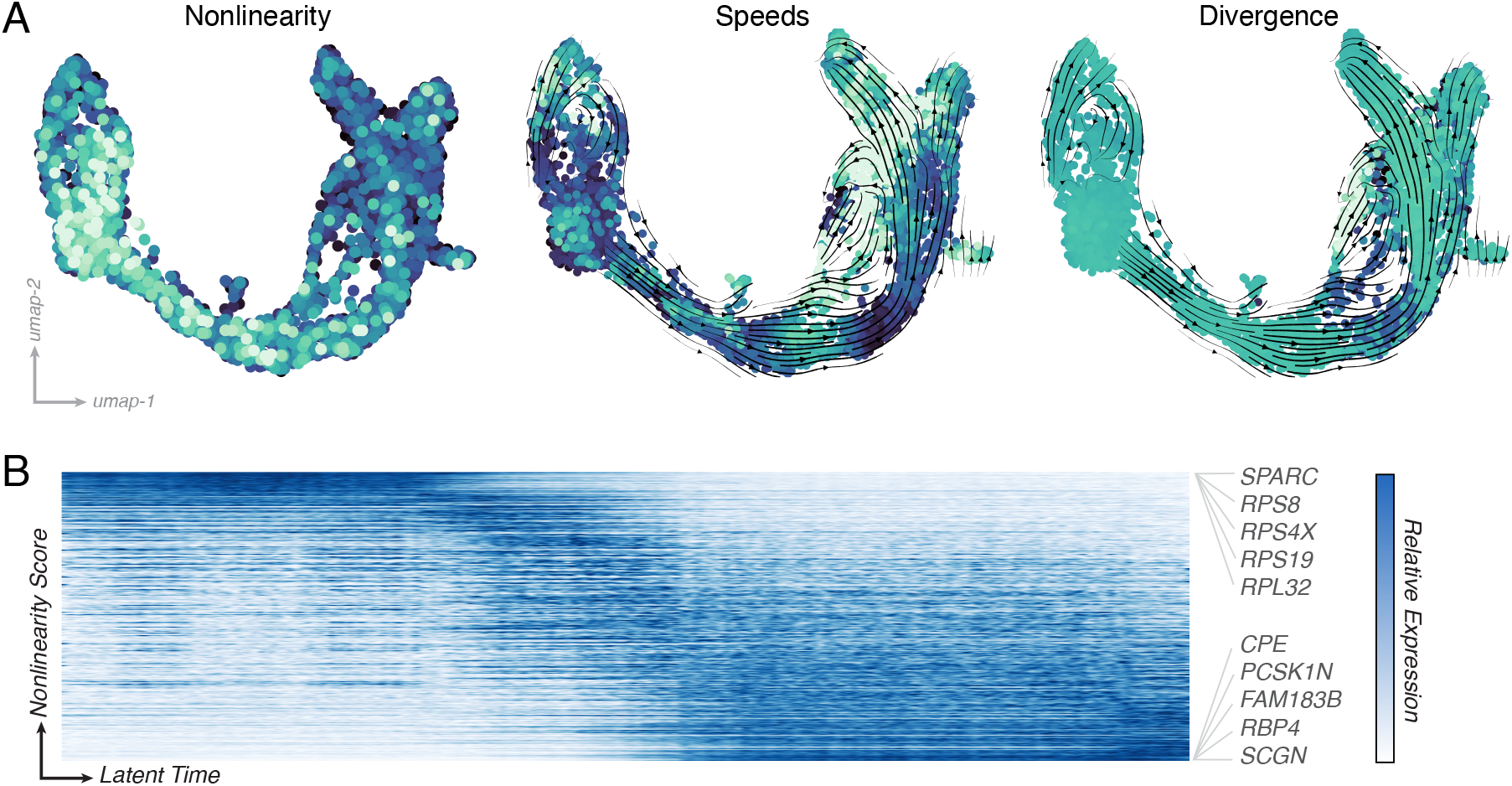
RiCE correlates with the speed and divergence of transcriptional dynamics. (A) The average nonlinearity, speed, and divergence of 3696 cells during endocrine development in the pancreas, during lineage commitment to four major fates. Dynamical metrics are computed using the RNA velocity for each cell and its nearest neighbors in the full expression space. (B) Expression levels versus the latent time inferred by RNA velocity. Genes are sorted by their correlation with the nonlinearity, and the highest and lowest-nonlinearity genes are annotated.

## III. DISCUSSION

We have introduced RiCE, a physics-based method for inferring causal relationships from time series. We show that many standard network inference approaches fail to extract physically-meaningful interactions when applied to dynamical systems, motivating the need for methods that identify mechanistic interactions. Our nonlinear causality-based approach leverages the unique properties of coupled dynamical systems, allowing it to achieve competitive performance with large-scale gene network inference methods, with its performance increasing as the degree of nonlinearity of the underlying system increases.

We find that our approach effectively reconstructs gene regulatory networks on known benchmarks, and that it finds meaningful dynamical signatures when applied to gene expression during known nonlinear biological processes like the epithelial-mesenchymal transition, endocrinogenesis, or immune cell differentiation. The primary limitation of our approach stems from its reliance on temporal or pseudotemporal data, limiting its scalability to large cell atlases containing multiple lineages. Recent work on local manifolds in high-dimensional dynamical systems offers routes for potential generalizations, including principled methods for decomposing large atlases into local dynamical systems associated with specific processes, each of which may be amenable to causal analysis [33–35]. More broadly, our approach establishes that nonlinear relationships present in many biological processes produce measurable signatures in experimental data. These cryptic interactions make appealing targets for further experimental studies quantifying the prevalence of nonlinearity as a general regulatory motif.

## Appendix A. CODE AVAILABILITY

A Python implementation of RiCE, including instructions, documentation, and the code for running benchmark experiments, is available at https://github.com/williamgilpin/rice

## Appendix B. STANDARD NETWORK INFERENCE ALGORITHMS FAIL TO DISCOVER NONLINEAR INTERACTIONS IN PHYSICAL SYSTEMS

To demonstrate a failure mode of common network inference strategies, we apply network inference approaches to time series generated by a set of strongly-coupled deterministic physical variables. We consider the Kuramoto model of coupled phase oscillators in a ring configuration,

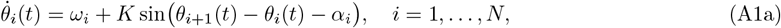

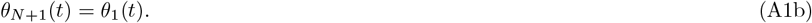

**Figure S1.**
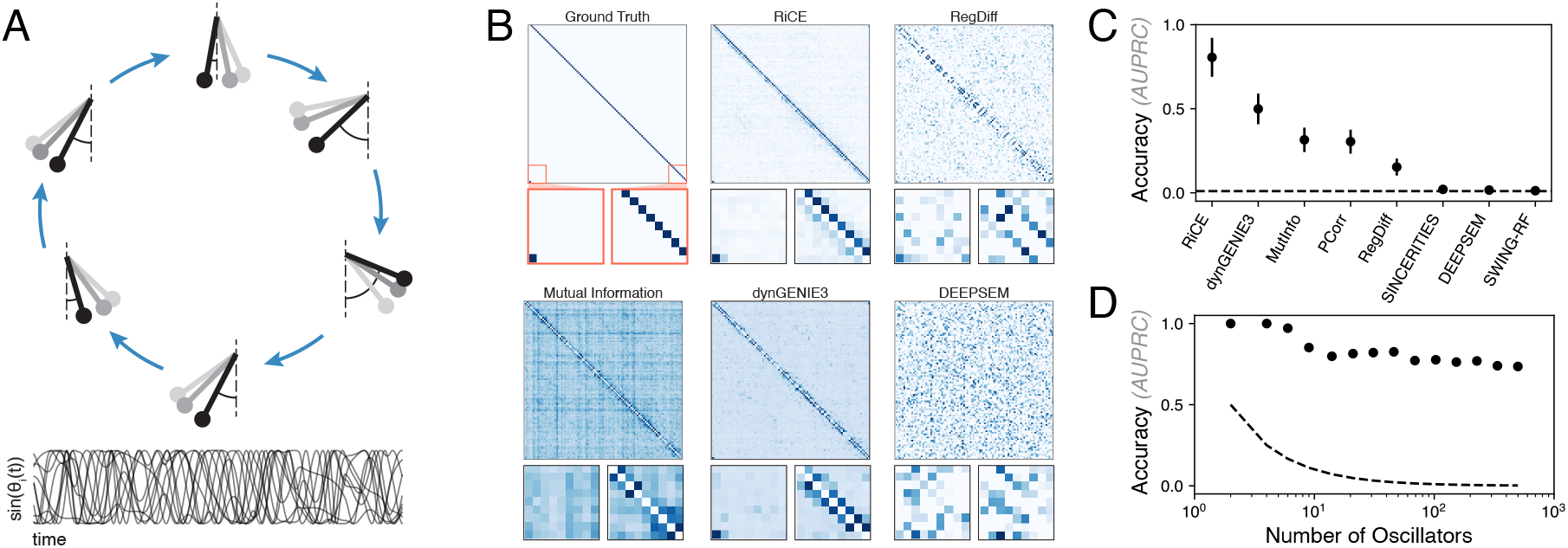
Detecting nonlinear interactions in coupled oscillator dynamics. (A). A schematic and example trajectories from a network of coupled phase oscillators. (B) The ground truth coupling matrix and the result of various network inference algorithms. Inset boxes highlight the periodic boundary condition and the off-diagonal non-reciprocal coupling elements. (C) Accuracy of different network inference algorithms. (D) Accuracy of RiCE as the number of oscillators increases. Horizontal line corresponds to the AUPRC of random guessing. Error bars are standard errors over 40 random initial conditions, sets of frequencies *ω*_*i*_, and phase offsets *α*_*i*_.

The random natural frequencies *ω*_*i*_ ∼ 𝒩(1, *σ*) represent a form of quenched disorder that prevent oscillators from becoming indistinguishable, while the uniformly-distributed frustration terms *α*_*i*_ *∼* 𝒰(0, 2*π*) prevent the oscillators from synchronizing (*θ*_*i*_ = *θ* for all *i*) even when the coupling strength *K* is large. Because of the angle-sum identity sin(*θ*_*j*_ − *θ*_*i*_) = sin *θ*_*i*_ cos *θ*_*j*_ + …, the dynamics in these coordinates have a leading-order intrinsic quadratic nonlinearity.

We generate a set of trajectories in the strongly-coupled limit *N* = 100, *σ* = 0.2, *K* = 1 and generate a dataset consisting of 10000 timepoints covering a duration of 400 rotations (in units of ⟨*ω*_*i*_⟩ = 1). To ensure the variables remain bounded, and to emphasize minimal quadratic nonlinearity, we use measurement coordinates sin(*θ*_*i*_(*t*)) instead of analyzing *θ*_*i*_(*t*) directly.

We apply a variety of existing network inference approaches to the 10000 × 100 dimensional coupled oscillators dataset. When evaluating all methods, we use representative methods from our benchmark experiments (Appendix G), and we use the same hyperparameter selection strategy. Among all methods, RiCE recovers the underlying network structure with much higher accuracy, doubling the performance of the second-best performing method, RegDiff. Importantly, among all methods tested, only RiCE correctly resolves the non-reciprocal (off-diagonal) structure of couplings, as well as the periodic boundary conditions that apply to the last oscillator. As a result, only RiCE recovers the true underlying physical dynamics, in the sense of identifying the key mechanistic components of Eq. A1.

Additionally, we find that RiCE has slow dropoff of accuracy even as the number of coupled variables increases. A two order of magnitude increase in the number of genes is required to halve the AUPRC (Fig. S1C). The dropoff is therefore shallower than the 1*/N* scaling expected for an approach that tests for each pairwise link independently (including the case of random guessing). We thus conclude that the ensemble step, in which all upstream variables are jointly weighted to produce a downstream forecast, confers scaling advantages relative to the pairwise approach used in many network inference approaches.

## Appendix C. THE RICE ALGORITHM FOR CAUSAL GRAPH RECONSTRUCTION

We seek to construct a matrix **c** ∈ ℝ^*M* ×*M*^ describing the causal relationships among a set of *M* univariate time series. A large value *c*_*ab*_ implies that the dynamical system associated with the time series indexed by *a* causally drives the dynamical system associated with the time series indexed by *b*, and so we use the notation *c*_*a*→*b*_ to emphasize this relationship.

### 1. Embedding and constructing the distance matrix

Given two univariate time series 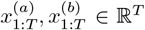, we perform a time-delay embedding with hyperparameter *d*_*E*_ to produce the lifted coordinates 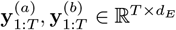.

For each timepoint in the lifted space, the *K* ≡ *d*_*E*_ + 1 nearest neighbors are found in the embedding space

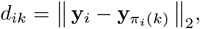

where *i* = 1, …, *T, k* = 1, …, *K*, and *d* ∈ ℝ^*T* ×*K*^. Here, *π*_*i*_(*k*) is the index of the *k*^*th*^ nearest-neighbor of a point **y**_*i*_ among the set {**y**_*j*_} _j ≠ *i*_

#### Explanation

This step invokes Takens’ theorem, a classical result in nonlinear dynamical systems that states that a finite number of time delays of a low-dimensional signal may be used to estimate the manifold of a higher-dimensional attractor that smoothly deforms onto the system’s true attractor. As a result, neighbor indices *π*_*i*_(*k*) encode information about the attractor topology, while the distance matrix *d* contains information about the attractor’s metric. Based on theoretical considerations, we set default hyperparameter values *d*_*E*_ = 3, the minimum dimension needed to embed aperiodic dynamics, and we set *K* = *d*_*E*_ + 1 = 4, corresponding to the minimum number of neighbors to triangulate a point.

### 2. Accounting for local attractor topology

For each lifted timepoint 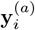, we use the distances *d*_*ik*_ and indices 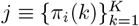 of its *K*-nearest neighbors to construct a self-forecast by fitting a kernel-weighted linear model,

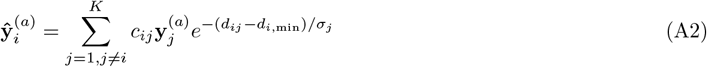

where *d*_*i*,min_ ≡ min_*k*_ *d*_*ik*_. There are two fitting parameters, the weighting matrix **c** ∈ ℝ^*T* ×*K*^ and the local metric ***σ*** ∈ ℝ^*T*^. We minimize the regularized mean squared reconstruction error 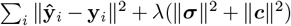, where *λ* is determined by cross-validation. In practice, we find that our results are consistent as long as *λ* is set to a high value, due to the large number of free parameters. We speed this calculation by solving first for the row-wise metric ***σ*** using nonlinear optimization, followed by ridge regression to determine the weights **c** of different neighbors [36].

#### Explanation

This step constructs a data-driven model of the upstream dynamical system associated with the variable *x*^(*a*)^. The model assumes that the dynamics evolve along a smooth attractor, and so a given point’s value may be approximated as a linear combination of its neighbors, weighted by their relative distance *d*_*ik*_ and the local flatness *σ*_*i*_.

This approach generalizes classical S-map analysis, a method for identifying global nonlinearity in a time series [8]. However, classical S-map assigns a single overall scalar nonlinearity score for the entire attractor, while our approach first assigns a separate nonlinearity score *σ*_*i*_ to each timepoint in a time series. For a smooth dynamical system, *σ*_*i*_ is related to the local flatness of the attractor, which is itself determined by the determinant of the instantaneous Jacobian stability matrix of the underlying dynamical system.

### 3. Creating a forecast

Using the fitted weight matrix and coefficients, we construct a forecast of the downstream variable,

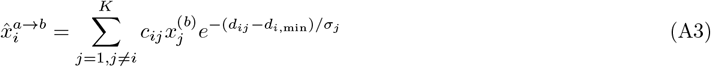

where all parameters *c*_*ij*_, *d*_*ij*_, *σ*_*j*_ correspond to the values found using the dynamical model in the previous step.

#### Explanation

The resulting prediction series 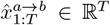 estimates the true downstream univariate time series 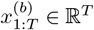 using the geometric model trained on the upstream time series 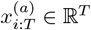.

### 4. Scoring a forecast

We next evaluate whether any linear combination of upstream variables predicts a given downstream variable. We perform Canonical Correlation Analysis (CCA) between the set of predictions produced by each upstream variable 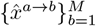, and a given downstream variable *y*^(*b*)^. In particular, we solve the ridge regression with constant regularization *λ* = 10^−3^,

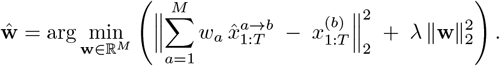

we then find the coefficient of determination,

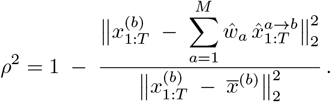

Where 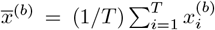. We also compute the *p*-value of this quantity assuming a *t*-distribution of *ρ* with sample size *M*. The values *ρ* and *p* are, respectively, the value and significance level of the Pearson correlation between the true downstream time series and its prediction using a linear combination of upstream variables. We use the correlation to scale the elements of the CCA assignment matrix **c** ∈ ℝ^*M*^,

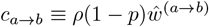

We iterate over all downstream variables *b*, and thus complete the causal matrix **c** ∈ ℝ^*M* ×*M*^ by iterating over columns (downstream variables).

#### Explanation

This step evaluates the quality of the forecasts 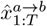 of the downstream time series 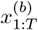. It evaluates whether any linear combination of the predictions 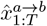 across all upstream time series *a* = 1, 2, …, *M* predicts the values of a single downstream time series 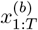. A high score therefore indicates whether the geometry of the dynamics of a given upstream variable indexed by *a* confers predictive information about a given downstream time series indexed by *b*.

### 5. Scaling for monotonicity

To improve the robustness of the RiCE score, we repeat the steps above for a modified dataset in which the number of timepoints is downsampled.

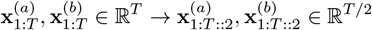

we refer to the recalculated causal matrix as **c**^*T*^ ∈ ℝ^*M* ×*M*^. We repeatedly downsampled the time series, and therefore sweep *T* over a logarithmic scale. We compute the Spearman correlation 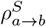 between each matrix element’s values 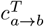 and the timescale *T*, and then rescale the values of the causal matrix

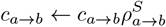

#### Explanation

This step sparsifies the causal matrix by discounting relationships among individual variable pairs *a, b* that do not improve with additional timepoints. In classical empirical dynamic modelling, a similar approach is used to eliminate spurious associations, which do not arise due to underlying dynamical relationships between pairs of variables [6].

### 6. False positive control (Optional)

While we do not require this setting in our main experiments, we include the optional ability to enforce a directed acyclic graph structure. If the causal score between a pair of variables *X, Z* is weaker than the score between either variable and an intermediate variable *c*_*x*→*z*_ *< c*_*x*→*y*_, *c*_*x*→*z*_ *< c*_*y*→*z*_, then the connection between *X* and *Z* is removed *c*_*x*→*z*_ → 0.

#### Explanation

Recent works show that network inference methods have a high rate of false positives, due in part to the phenomenon of *causal transitivity*, in which pairwise causal detection algorithms discover indirect interactions [37, 38]. This effect is pronounced in gene regulatory networks with low amounts of self-regulation, in which case the underlying network may have a bipartite structure consisting of transcription factors and their downstream targets. We therefore optionally prune an inferred network *c*_*a*→*b*_ into a directed acyclic graph.

Our approach is motivated by a similar calculation based on the data-processing inequality, that has previously been used to prune gene regulatory networks inferred using information-theoretic methods [39]. For coupled dissipative systems, a similar relationship holds due to the relationship between the sub-Lyapunov exponents of coupled systems and information transfer between partially-synchronized dynamical systems [40–42]. Because the mutual information between two coupled systems is bounded by their degree of synchronization, a violation of the data processing inequality may be used to test for indirect synchronization.

## Appendix D. RELATED WORK AND LIMITATIONS

### Prior Benchmark Studies

Diverse studies consider the DREAM4 time series benchmark, with recent works finding that purely statistical approaches (which do not directly use temporal information) achieve accuracy scores matching leading time series approaches [1, 43]. However, the small size of this dataset, and plateauing scores, suggest a degree of saturation on this dataset. A benchmark of methods, including statistical approaches like ARACNE and CLR and dynamics-based methods like SCODE, found that no approach consistently performed well, with different methods exhibiting wide variance in performance across different simulated and experimental single-cell RNA sequencing datasets [44]. A study introducing the BEELINE benchmark of single-cell pseudotime data found no substantial advantage in dynamics based models over statistical models [10], a result affirmed by later work showing that time-agnostic probabilistic machine learning models perform state-of-the-art on this dataset [45, 46]. Later work on simulated single-cell datasets found that dynamics-based approaches only outperform classical statistical approaches on linear trajectory datasets, and not bifurcating datasets, particularly in low-noise settings [47]. This observation is consistent with pseudotime structure limiting the effectiveness of current dynamics-based methods. A newly-introduced single-cell gene expression benchmark, McCalla, compares eleven computational methods for inferring gene regulatory networks from single-cell RNA-seq across seven datasets in human, mouse, and yeast [7], This study profiles runtime and memory and excludes non-scalable methods (SCHiRM, HurdleNormal, BTR). Benchmarked algorithms span from information-theoretic (PIDC, Scribe), tree-based regression (SCENIC), ODE-based (SCODE), Gaussian graphical models (SILGGM), and a Pearson correlation baseline. Gold standards consist of ChIP-seq, perturbation, and a combined gold standard. Methods were scored using AUPRC, F-score, and predictable TF metrics. Although global recovery was modest, local metrics revealed consistent identification of key regulators (e.g., POU5F1, SOX2, NANOG in ESC differentiation, or NF-*κ*B in dendritic cells). Imputation via MAGIC seldom improved accuracy, and scRNA-seq-derived networks performed on par with bulk-derived networks. Incorporation of motif-derived priors and NCA-based transcription factor activity estimates in MERLIN and Inferelator yielded the greatest performance gains. A recent study uses the McCalla dataset to show that newer methods, including neural-network based generative models DEEPSEM (discussed below) and scPrint, a pretrained foundation model, achieve leading scores on this dataset [48].

### Empirical dynamic modelling

RiCE conceptually generalizes convergent cross-mapping (CCM), a data-driven method that infers causality by determining whether geometric information in one time series may be used to successfully predict another time series [6]. While CCM has proven widely-applicable in data-limited fields like ecology and environmental science, it has seen less application in gene expression, where small dataset sizes and large number of genes can lead to high false-positive rates. Like CCM, our approach consists of using time-delay embeddings to estimate dynamical manifolds, and then using forecast models trained on one time series to predict another. Unlike CCM, we fit an adaptive local model of the dynamics using approaches drawn from topological data analysis, producing an instantaneous nonlinearity score for each timepoint and gene based on an estimate of the local stability matrix of the underlying dynamical system (Appendix C). Further differing from CCM, we construct forecasts using an ensemble of forecast models separately trained on each upstream driver, rather than isolating each pair of time series. Unlike CCM, our forecast models for each upstream variable consist of kernel regressions over neighbors, rather than sums over neighbors weighted by their distance in embedding space. This last component resembles S-map, another concept from the forecasting literature; however, our kernels are adaptive based on the local stability matrix, rather than based on setting a single global kernel hyperparameter [8]. Our approach differs from multiview embedding, which separately performs neighbor identification and simplex forecast model training for each potential upstream time series, and then uses the best *k* upstream models to forecast the downstream time series (using traditional simplex forecasting methods) [49]. Unlike partial cross mapping, we eliminate indirect interactions through post-hoc pruning, rather than conditioning the forecast accuracy calculation on the indirect variable [50]. Additional technical distinctions differentiate our approach from other nonlinear causality approaches, particularly our use of fast approximate nearest neighbor search, adaptive neighbor weighting using an estimate of the local stability matrix, exponential striding of time series to prune for improvement with time series granularity, and optional pruning for indirect interactions.

### Strong versus weak causality

Our observational approach detects weak causality, defined as a relationship among two variables in which one variable dynamically forces a second dynamical variable. For example, a strong relationship *c*_*a*→*b*_ may imply a skew-product dynamical system of the form 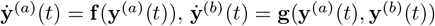. Weak causality therefore implies a deterministic relationship as well as a predictive mechanism, in line with causality typically considered in studies of synchronization in dynamical systems [42, 51–54]. Our causal definition therefore differs from stronger forms of causality established through interventional studies or by directly testing counterfactuals [55]. Such approaches require access to the underlying generator of the data, such as the experimental system or simulation model.

### Other observational causality methods

Nonlinear causality methods like RiCE and CCM resemble methods like Granger causality because they use information from covariate upstream time series to construct joint forecasts of downstream time series. However, unlike Granger causality, nonlinear methods seek to address (1) non-separable linear systems, (2) weakly-coupled variables, and (3) disentangle shared confounding driving variables from direct interactions [6, 56]. Direct generalizations of Granger Causality, such as Neural Granger Causality, seek to address the first limitation only, by replacing the regularized linear regression model typically used to create a joint forecast in traditional Granger Causality [57].

**Figure S2.**
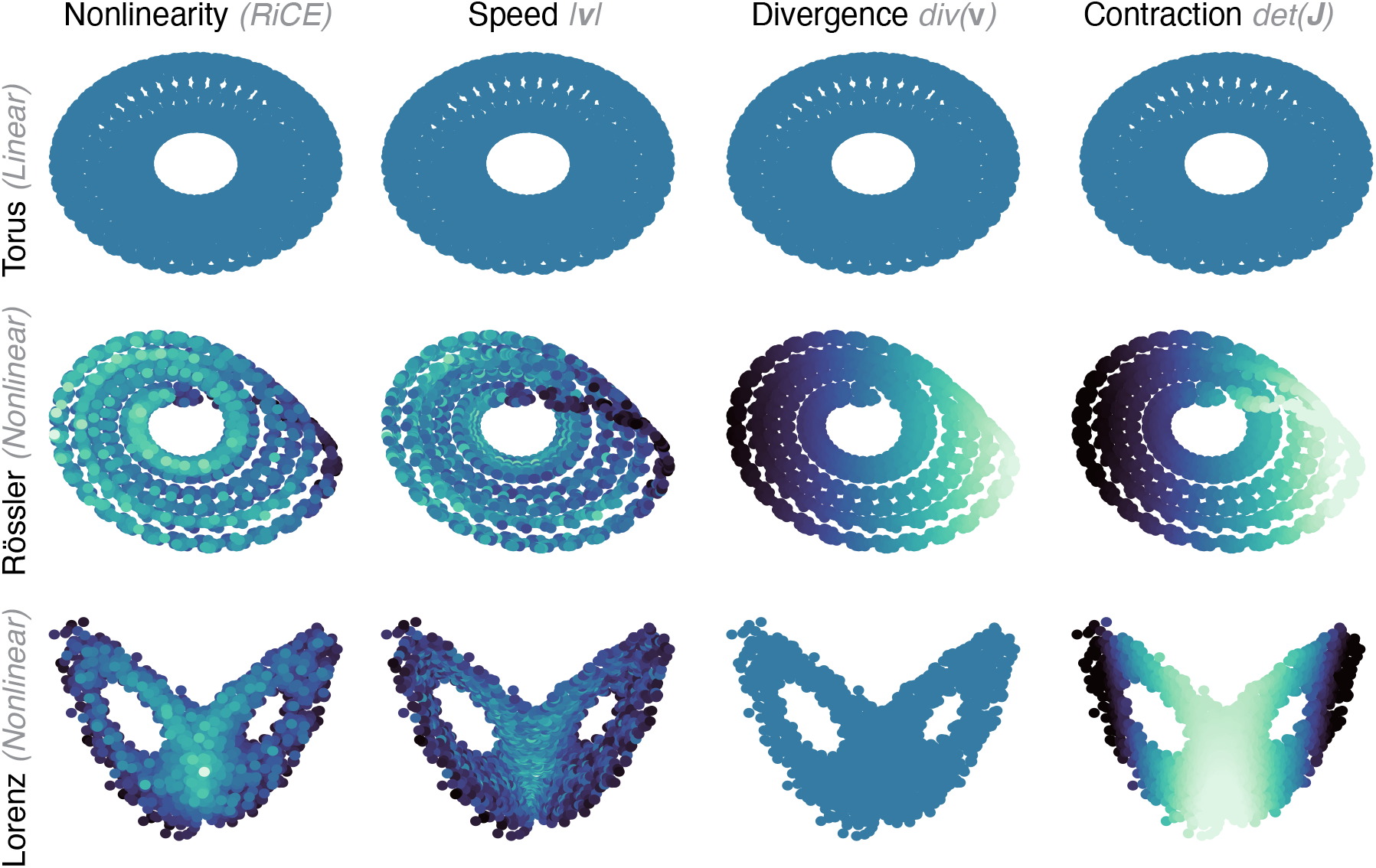
Local properties of nonlinear dynamical systems. The RiCE nonlinearity score is computed within RiCE purely from the time series. The speed, divergence, and Jacobian determinant all use different instantaneous properties of the underlying dynamical equations. For the case of the Lorenz attractor, the divergence is a constant negative value everywhere on the attractor. For a linear, bounded dynamical system like the torus, all quantities are constant.

In longitudinal studies, the number of distinct observable timepoints may be limited. Spatial CCM overcomes the limitations of short time series when multiple nearly-replicated short time series are available—such as simultaneous measurements of similar subjects or nearby locations. These time series are then aggregated, and points are sampled from this pool when constructing simplex forecasts [58]. Partial cross-mapping addresses causal transitivity by using partial correlations to control for shared drivers [50].

## Appendix E. PROPERTIES OF NONLINEAR DYNAMICAL SYSTEMS

We review general properties of nonlinear dynamical systems, and explore how the RiCE nonlinearity score calculated internally within RiCE reveals local information about the system’s dynamics.

### Linear dynamical systems

Consider the autonomous linear dynamical system in *N* dimensions,

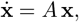

where *A* ∈ ℝ^*N* ×*N*^ and **x** ∈ ℝ^*N*^ The Jacobian matrix is constant:

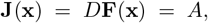

hence

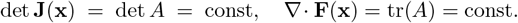

**Figure S3.**
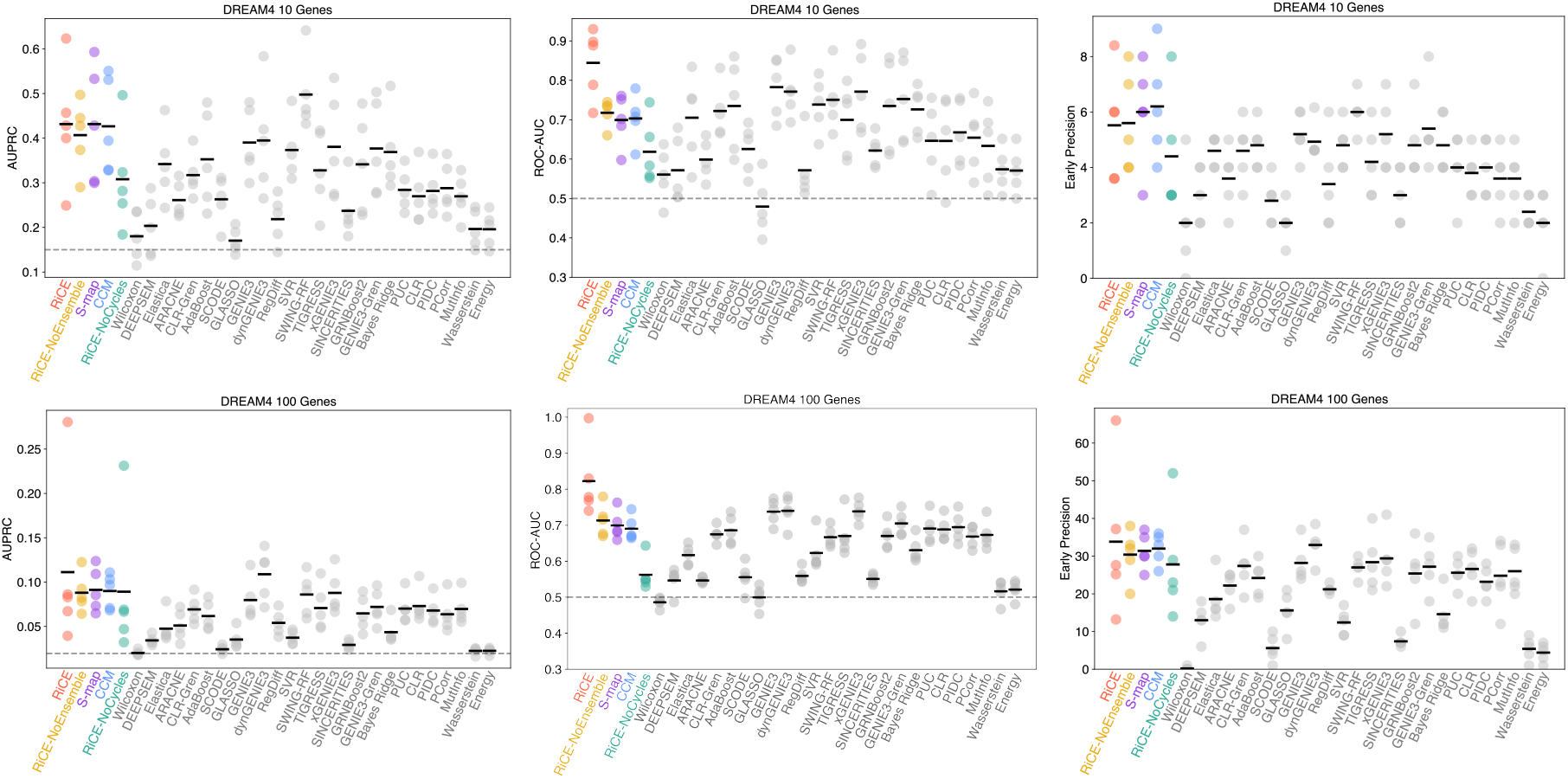
All metrics on the DREAM4 challenge. Accuracy of network link identification (AUPRC, AUROC, and Early Precision) for the 10 gene (top row) and 100 gene `InSilico` Challenge. Method names and ablations match the descriptions in the main text. Dashed lines correspond to the expected values for a random classifier.

In order for the dynamics of a linear dynamical system to remain bounded, the state of the system must either approach a fixed point or limit cycle. In the former case, all dynamical measurements necessarily become constant. For the case of a limit cycle, *A* is skew-symmetric (*A*^T^ = −*A*), and so *e*^*At*^ is orthogonal and the flow is bounded with

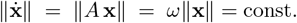

### Nonlinear dynamical systems

For a general autonomous nonlinear system

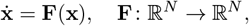

the Jacobian matrix

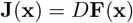

varies with **x**. If the system exhibits an extended attractor (limit cycle or strange attractor), then the Jacobian varies with *t* even at long times. Consequently, the fields

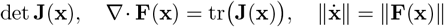

are in general non-constant on the state space and serve as local probes of the system’s instability and nonlinearity. The *Speed field s*(**x**) = ∥**F**(**x**)∥ measures the local rate at which trajectories traverse the attractor. Regions of large *s*(**x**) correspond to rapid motion and potential local stretching. The *Divergence* ∇·**F**(**x**) of the flow field quantifies infinitesimal volume contraction (*<* 0) or expansion (*>* 0), directly relating to the local stability or instability of nearby trajectories. The *Jacobian determinant* det **J**(**x**) gives the oriented volume–change factor for an infinitesimal neighborhood under the flow map, highlighting folding and stretching due to nonlinearity.

We show the properties of these metrics in Figure S2 for two exemplary nonlinear systems with bounded attractors, as well as the same calculation performed on a quasiperiodic linear system.

## Appendix F. SCALING PROPERTIES OF RICE

In Figure S5A, we compare the theoretical and observed scaling of RiCE as the number of cells (timepoints) and genes (features) increases, and find agreement.

**Figure S4.**
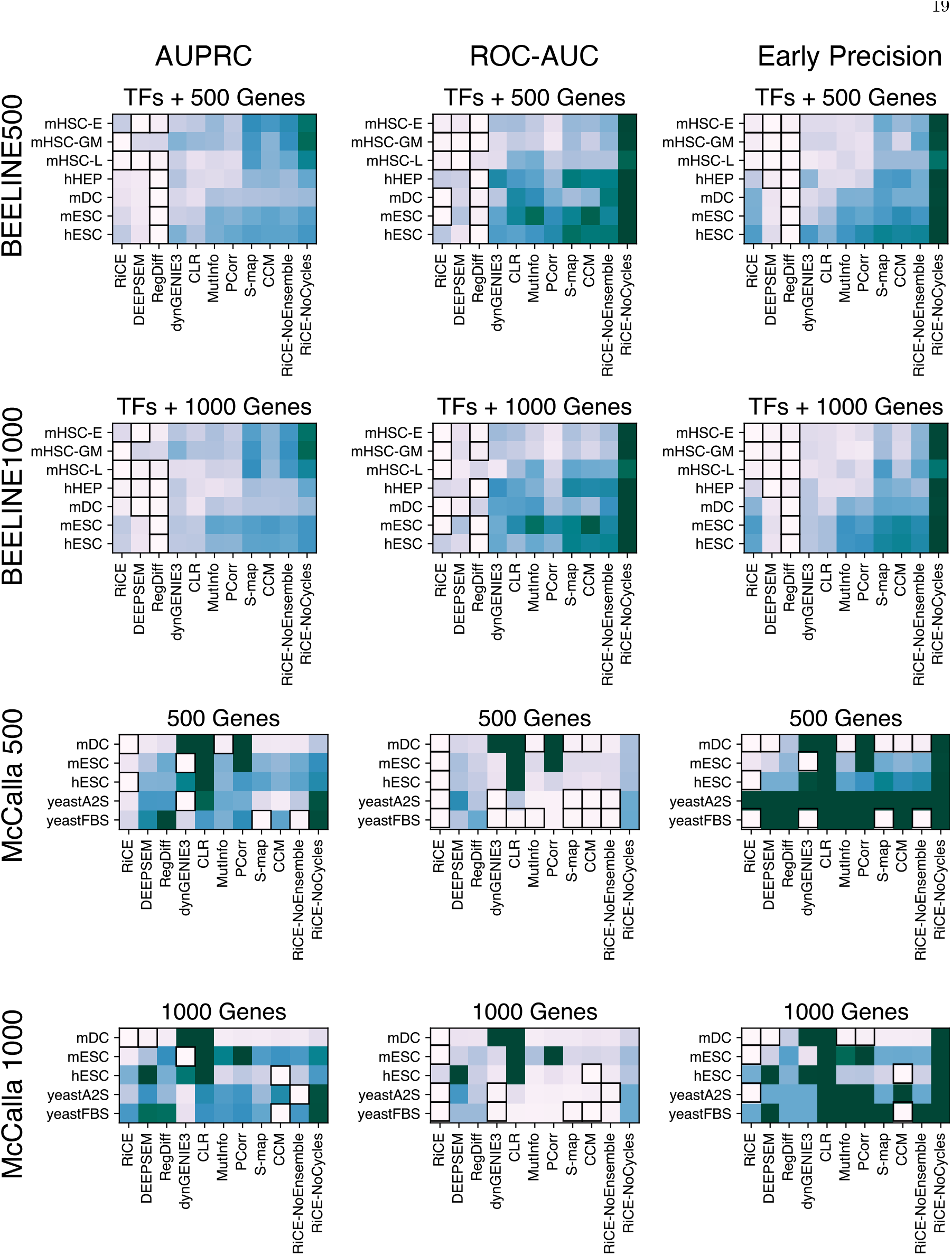
All metrics on the BEELINE and McCalla datasets. Boxes indicate highest-performing methods for each dataset within one standard error across bootstrapped replicates (resampled cells).

**Figure S5.**
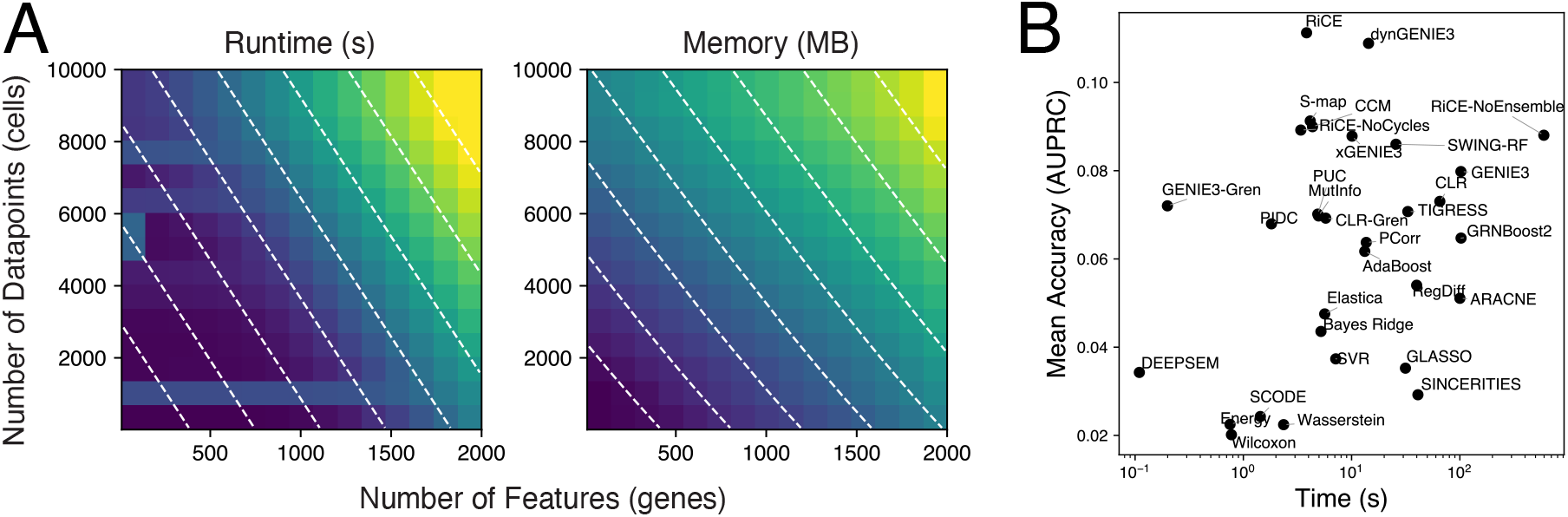
Runtime and memory scaling versus performance. (A) The runtime and maximum memory used by RiCE when running on a single processor. Dashed lines correspond to evenly-spaced isocontours of the theoretical expected scaling of 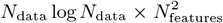 for runtime and 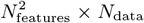 for memory. (B) AUPRC of each method on the DREAM4 gene challenge, versus its runtime on a single CPU. We note that several methods, like GENIE3 and its variants, can be parallelized, while other methods like RegDiff and DEEPSEM, run much faster on GPU.

### Runtime

The runtime of RiCE with the number of genes *M* (time series features) is set by the requirement to build the full causal graph among all pairs. The runtime therefore grows as ~ 𝒪(*M*^2^). Regarding the number of timepoints, cells, or datapoints *N*, the runtime is set by a combination of the penalized linear regression (∼ 𝒪(*N*)) and the repetition of the algorithm across different partitions (∼ 𝒪(log *N*)). The overall runtime scaling is therefore ∼ 𝒪(*N* log *NM* ^2^).

#### Memory

The peak memory usage of RiCE with the number of genes *M* (time series features) is also set by the requirement to build the full causal graph among all pairs (∼ 𝒪(*M* ^2^)). The memory bottleneck with *N* is also the linear regression step, (∼ 𝒪(*N*)). The overall memory scaling is therefore ∼ 𝒪(*NM* ^2^).

## Appendix G. BENCHMARK EXPERIMENT DESIGN

### A. Datasets

We consider 5 distinct datasets, including 2 newly-developed by us, in order to evaluate the generality and robustness of our approach relative to others.

### Smoketest

We create a trivial dataset in which each time series *X*^(*k*)^ consists of univariate Gaussian noise, 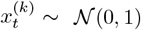. We induce artificial correlations among variables by randomly sampling 1*/N* pairs of nonidentical indices, *ℓ, m, ℓ ≠ m*, and then fusing the corresponding time series via the update rule *X*^(*ℓ*)^ ← (1 − *β*)*X*^(*m*)^ + *β X*^(*ℓ*)^. For our dataset, we set *N* = 20 and continuously vary *β* over 15 evenly-spaced values between 0.0 and 1.0.

This dataset serves to evaluate whether the benchmark models and metrics are correctly implemented. The best possible performance on this dataset can be achieved by taking the Pearson correlation among all pairs of time series, and using the resulting matrix as a reconstruction of the regulatory network. As *β* varies, the AUPRC performance of this method (termed PCorr, or partial correlations, in our experiments) varies between 1*/N* (when *β* = 0, there is no meaningful correlation structure besides random chance) and 1 (when *β* = 1, some time series are exactly duplicated in the dataset).

We find that most benchmark methods exhibit AUPRC proportional to *β* (Fig. S6), indicating that they detect no meaningful links when little information is available, and their accuracy improves as correlation structure becomes more apparent. All methods perform better than random guessing when *β >* 0, assuring that our benchmark implementations and hyperparameters extract meaningful information. Several methods perform better than random chance even when no meaningful structure exists in the dataset (*β* = 0), indicating a systematic bias towards amplifying weak correlations—a potential result of the small size of our dataset. We find that the primary cause of variation in performance across methods stems from the degree to which different methods can exploit statistical correlation structure in the data. The statistical and information-theoretic approaches perform more strongly, while methods specialized to exploit temporal information, like SWING-RF or dynGENIE3, seemingly underperform because they have comparatively less advantage on this non-temporal dataset.

**Figure S6.**
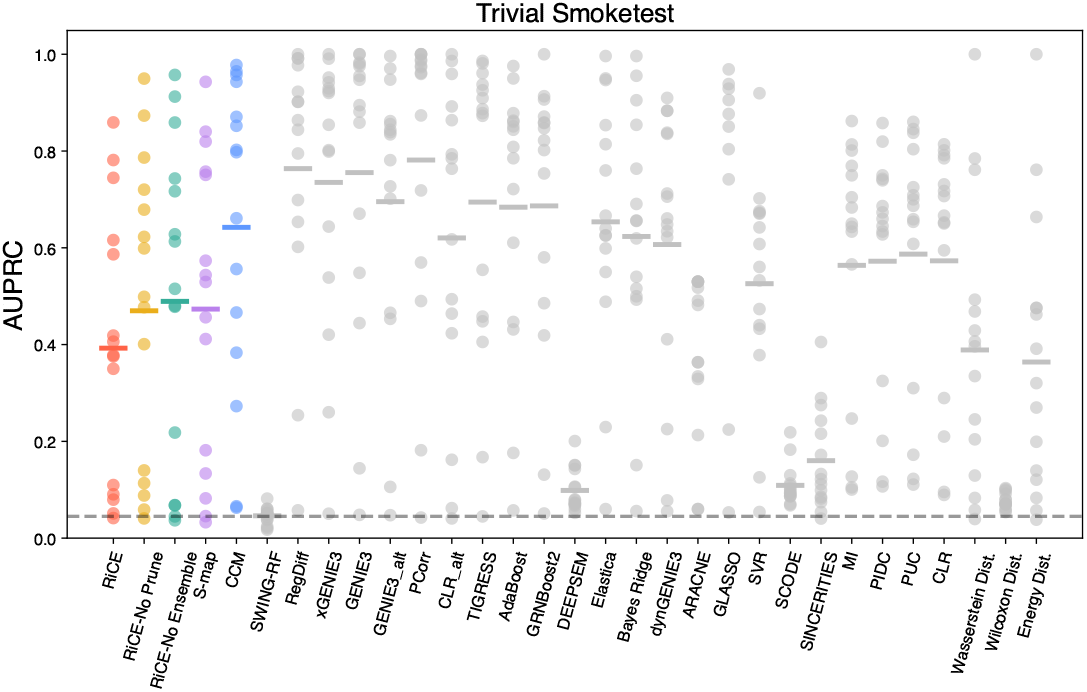
Performance of benchmark models on a trivial statistical dataset. Accuracy of network link identification (AUPRC) on datasets in which pairs of time series exhibit direct statistical correlations. Vertical points correspond to replicates in which the strength of the specified correlations varies from 0 (fully-independent time series) to 1 (duplicated time series present). The mean for each set of replicates is indicated with a horizontal line. The expected AUPRC of a random classifier is indicated by a dashed line on the graphs.

### DREAM4

The DREAM4 benchmark represents stochastic reaction kinetics for 10 networks consisting of 10 or 100 genes [9]. Each network represents a synthetic gene regulatory system with topology drawn from biologically-plausible submodules sampled from experimental transcriptional interaction maps from *E. coli* and *S. cerevisiae*, ensuring that connectivity patterns (e.g. degree distributions, motif frequencies) reflect those observed in real organisms. Dynamics are simulated with stochastic differential equations (incorporating intrinsic noise and regulatory kinetics) using the GeneNetWeaver framework, which implements Hill-type (or Michaelis–Menten) reaction kinetics to govern transcription and degradation processes [59]. For each 10-gene network, five microarray-style time-series (21 time points each) and multifactorial steady-state data under mild basal-transcription perturbations are generated; for each 100-gene network, ten time-series replicates are produced. Static perturbations are applied during the first half of each series to drive adaptation, followed by relaxation phases, and systematic knockout/knockdown datasets are provided to emulate genetic perturbations across every gene.

### Twist

We introduce a new dataset that adds nonlinear dynamics to subnetworks resembling the ones originally used in the DREAM challenge. We load a reference network, and select an initial “seed” node at random, with selection probabilities proportional to node degree. We then iteratively expand the subnetwork outwards by sampling one neighbor of the current node weighted by neighbor degree, and adding both the node and the connecting edge if not already present. New nodes are enqueued either at the front (mimicking depth-first expansion) or the back (breadth-first) of the exploration list according to a probabilistic bias, and this process continues until the subnetwork reaches the requested size. If a directed subnetwork is requested, a random half of its edges are reversed at the end to introduce additional structural variation. We also include the original DREAM4 networks. Following DREAM4, as the *E. coli* reference network we use RegulonDB v6.7, which aggregates transcription-factor–target links manually curated from biochemical footprinting, binding assays and motif analyses drawn from the literature [60]. For *Saccharomyces cerevisiae* we use a network built by integrating genome-wide TF–DNA binding evidence (chiefly ChIP–chip) with computational motif scans and literature-curated interactions [61].

On each network, we simulate expression dynamics using a mechanistic dynamical model of coupled mRNA and protein dynamics [62]. This model captures key features of genetic regulatory systems: autoregulatory positive and negative feedback, dimerization of transcription factors, and stimulus-dependent phosphorylation, and as a result this model exhibits diverse dynamics including multistability, oscillations, and frequency-selective tuning. We use a simplified version of the model that reduces explicit mRNA kinetics and phosphorylation-state variables, reducing the dynamics to a discrete map with improved stability in discrete time [63].

### BEELINE

A set of curated single-cell RNA sequencing datasets consisting of mouse and human development and proliferation processes [10]. Each cell has been annotated with pseudotime via SLINGSHOT, an algorithm that constructs a minimum spanning tree among cells and then assigns pseudotime via principal curves [64]. Following prior works, we consider two variants of the dataset: (1) the top-500 highest variance genes and all experimentally-tested transcription factors, and (2) the top-1000 highest variance genes and all experimentally-tested transcription factors. BEELINE includes multiple experimental estimates of ground-truth regulatory methods, including ChIP-seq (both nonspecific and cell-type specific), protein interaction networks (STRING), and loss-of-function/gain-of-function perturbation experiments. Prior works report highest scores when using STRING as the ground truth [10]; moreover, this reference contains the highest coverage of interactions. We thus select STRING as our ground-truth networks.

### McCalla

A similar single-cell RNA sequencing dataset to BEELINE, with newer and larger datasets [7]. Five publicly available scRNA-seq datasets are curated, spanning three species: two yeast stress response experiments (yeastA2S, yeastFBS), one human embryonic stem cell differentiation time course (hESC), one mouse dendritic cell response (mDC), a mouse reprogramming dataset (mESC). Cell counts range from 163 to 36199 cells. Each dataset is filtered to retain only genes detected (count *>* 0) in at least 50 cells and cells with ≥ 2000 total UMIs; known transcription factors and signaling proteins were re-included if expressed; count matrices were depth-normalized and square-root transformed. For each species and cell state, three “ground truth” networks are compiled: (1) ChIP-derived interactions, (2) regulator perturbation (knockdown/knockout) interactions, and (3) their intersection. In order to apply dynamics-based analysis to these networks, we calculate pseudotime using diffusion components [12].

## Appendix H. BENCHMARK DESIGN

We do not use external annotations (like transcription factors) or prior information about the network, and instead only evaluate models’ ability to extract information from time series. For this reason, we report lower scores on DREAM4 than methods that are given transcription factor identities.

### A. Metrics

We primarily use the AUPRC due to its common use and ability to correct for the sparsity of gene networks. However, we also consider AUROC and Early Precision, two other informative measures of network reconstruction quality [10]. Given a network inferred among *N* genes, the AUPRC and AUROC are calculated over the predictions for all *N* (*N* − 1) predicted links. The baseline scores (score of a random estimator) for AUPRC is the network density divided by *N* (*N* − 1); for AUROC the baseline score is 0.5, and for Early Precision the baseline is the same as for AUPRC. We note that several benchmark studies that subselect known transcription factors score their models only on this list of known targets, producing higher AUPRC scores than we report.

## Appendix I. BENCHMARK MODELS

We select benchmark methods based on their ability to operate in a fully-blind setting (without any annotations of transcription factors), the availability of documented implementations (ideally by the original authors), their frequency of usage, and the diversity of their approach. When possible, we use each method’s authors’ original code, and we do not tune user-set hyperparameters beyond their default values (including for our own method). The performance of a given method may vary depending on initialization and hyperparameter settings, and so we avoid extensive tuning. We do not provide any models with prior estimates of transcription factors or ground-truth links. To ensure consistency in benchmark evaluations, for any methods that accept transcription factor candidates, we provide a list of all *N* genes. We include several methods from the GReNaDIne package, which provides reference implementations of many network inference approaches [65], this leads to two distinct implementations of GENIE3 and CLR among our benchmarks. We group the benchmark models based on the type of network inference approach.

### A. Statistical Models

#### Partial correlations (PCorr)

Linear correlations in expression calculated across all cells or timepoints [66, 67], which can be interpreted as measuring joint expression probabilities across the population.

#### ARACNE

Mutual information between expression profiles with surrogate data testing [39, 68]. This method uses the data processing inequality, a concept from information theory, in order to prune the causal graph of spurious links arising from indirect interactions, a phenomenon known as *causal transitivity*.

#### Mutual Information (MutInfo)

The empirical mutual information between two genes, averaged across all time-points [69, 70].

#### Context Likelihood of Relatedness (CLR)

The cumulative empirical distributions of mutual information values between two genes, relative to all other genes [69, 70].

#### Partial Information Decomposition (PIDC)

A multivariate generalization of mutual information that account for higher-order causal transitivity that may arise in single-cell datasets [71].

#### Graphical Lasso (GLASSO)

A sparse maximum likelihood estimator for the inverse of the covariance matrix of a multivariate Gaussian distribution describing an observed multivariate dataset. We set the L1 penalty hyperparameter by performing cross-validation over 5 logarithmically-spaced values between 10^−6^ and 10^2^.

#### xGENIE3

An adaptation of GENIE3 that replaces the random forest with an Extra Trees regressor to predict each gene’s expression from all others, leveraging the increased accuracy and regularization of gradient boosting; feature importances are used to infer regulatory links [72].

#### GENIE3

A random forest that predicts the expression profile of a given gene across the population, as a function of the other genes. Feature importance is then used to identify which genes are most informative for the regression, indicating a link between the genes [73].

#### GRNBoost2

A high-performance implementation of GENIE3 using gradient boosting machines optimized for single-cell data, which parallelizes tree building and employs early stopping to scale to tens of thousands of cells [74, 75].

#### Proportional Unique Contribution (PUC)

Decomposes mutual information in triplets of genes to quantify the unique pairwise information shared between two genes after accounting for higher-order transitive effects, improving specificity in single-cell network inference [71].

#### B. Regression Models

##### Trustful Inference of Gene REgulation using Stability Selection (TIGRESS)

Applies stability selection to LARS-based Lasso regression: for each target gene, LARS selects regulators in repeated subsamples, and selection frequencies yield a robust ranking of candidate edges [76].

##### Elastica

Employs the elastic net (a combination of Lasso and Ridge penalties) to regress each gene on all others, balancing sparsity and coefficient shrinkage to infer networks that are robust to multicollinearity [77].

##### Bayesian Ridge

A Bayesian linear regression with an automatic relevance determination prior (Gaussian prior on weights), yielding a sparse connectivity matrix via posterior mean weights and accounts for uncertainty in parameter estimates.

##### SVR

Applies support vector regression with a radial basis function kernel to model nonlinear gene-gene dependencies; the learned support vectors and regression coefficients quantify interaction strengths.

##### AdaBoost

Utilizes adaptive boosting to infer regulatory links: weak gene-gene regressors are trained iteratively, reweighting samples to focus on poorly predicted expression values, and the final ensemble’s feature weights indicate edge strengths [78].

### C. Physics-based Models

#### dynGENIE3

An extension of GENIE3 that uses a random forest to estimate a nonlinear forcing term in an underlying differential equation [79].

#### SCODE

Fits a system of linear ordinary differential equations to single-cell expression trajectories by projecting the high-dimensional dynamics onto a low-dimensional latent space, estimating a time-dependent regulatory matrix via least-squares under sparsity constraints [80].

#### Sliding-Window Inference with Networks and Granger causality, using random forests (SWING-RF)

Learns nonlinear autoregressive relationships within moving time windows, thereby capturing both instantaneous and time-lagged regulatory interactions [81].

#### SINCERITIES

Quantifies temporal shifts in single-cell gene expression distributions—using the Kolmogorov–Smirnov distance normalized by uneven time intervals—and fits a non-negative ridge regression (ridge parameter chosen by LOOCV) to predict each gene’s distributional change from those of all regulators; directed edges are ranked by regression coefficients and signed via Spearman partial correlations between expression profiles across all timepoints [82].

### D. Distances

#### Energy

Computes the distance correlation (energy statistic) between gene expression vectors to capture both linear and nonlinear associations; strong distance correlations imply regulatory links [83].

#### Wasserstein

The first-Wasserstein (Earth Mover’s) distance between the empirical expression distributions of two genes across cells or timepoints; smaller distances indicate similar distributional shapes suggestive of co-regulation [84].

#### Wilcoxon

Uses the Wilcoxon rank-sum test to assess whether the expression levels of one gene stochastically dominate those of another across the cell population; significant rank-based differences imply directed regulatory influence under the assumption that up-regulation follows activation.

### E. Artificial neural networks

#### Regularized Diffusion (RegDiff)

A diffusion probabilistic model that infers GRNs by (1) adding Gaussian noise to single-cell expression vectors according to a fixed diffusion schedule, (2) embedding noisy expression, time step, and cell-type features through stacked MLP blocks, and (3) learning a trainable adjacency matrix within the reverse denoising network whose weights directly quantify regulatory edge strengths; this approach avoids costly matrix inversions, allowing the code to scale favorably with large numbers of genes [46].

#### DEEPSEM

A beta-variational autoencoder that embeds each gene’s expression via shared MLPs, and explicitly models their interactions. The model encodes the adjacency matrix (mapping latent features to expression) and its inverse (decoding noisy observations), such that the learned encoder/decoder weights directly correspond to a sparse regulatory network. This generative model is trained end-to-end to jointly infer GRNs, generate low-dimensional embeddings, and simulate realistic scRNA-seq profiles [45]. Following the original work, we average the interaction matrices produced by 10 replicates with different random initializations, and we run the method in a manner that is not specific to cell type.

### F. Ablations

We remove each component of the RiCE algorithm, in order to evaluate their relative contributions to the overall algorithm’s performance:

**NoCycles** searches the discovered causal matrix for indirect interactions of the form *c*_*i*→*k*_ *< c*_*i*→*j*_, *c*_*j*→*k*_ prunes potential transitive interactions to produce a network with a traditional top-down directed regulatory structure.

**NoEnsemble** fits a separate forecast model to each pair of genes and thus removes any causal driving arising from combinations of upstream variables.

**S-map** removes both ensembling and adaptive reweighting of nonlinear neighbors, corresponding to a classical approach that fits a single, global nonlinearity parameter to each dynamical system [8].

**Convergent Cross-Mapping (CCM)** removes ensembling, adaptive weighting, and replaces all least-squares fits with distance-weighted averages over neighbors, corresponding to an earlier nonlinear causality method [6].

## Notes

https://github.com/williamgilpin/rice

### Competing Interest Statement

The authors are affiliated with Medici Therapeutics in Cambridge, MA. All code, results, and data discussed in the manuscript are available at the provided GitHub link.

